# M-ECG: Extracting Heart Signals with a Novel Computational Analysis of Magnetoencephalography Data

**DOI:** 10.1101/2025.04.17.649325

**Authors:** Aqil Izadysadr, Hamideh Sadat Bagherzadeh, Jennifer R. Stapleton-Kotloski, Gautam Popli, Cormac A. O’Donovan, Dwayne W. Godwin

## Abstract

Magnetoencephalography (MEG) measures the magnetic fields generated by neural activity with high temporal and spatial resolution. Because of its focus on brain activity, other biopotentials, including muscle artifacts and heart signals, are typically filtered or rejected. In this study, the feasibility of extracting cardiac signals from MEG data, which is termed “magnetoencephalographic electrocardiogram” (M-ECG; in contrast to the electrocardiogram or ECG) is explored. Using the publicly available Brainstorm MEG auditory dataset – CTF and OMEGA resting-state sample dataset, a novel algorithm is developed that utilizes either independent component analysis (ICA) or MEG reference sensors to extract M-ECG signals and compute heart rate variability (HRV) from MEG data reliably and accurately. Signal processing methods in the time, frequency, and time-frequency domains along with statistical tests such as Spearman’s correlation, root mean square error, mean absolute error, Bland-Altman mean difference, and Mann-Whitney U Test are employed to assess the similarities across the signals. The results indicate a significant alignment of temporal and frequency spectral power characteristics between M-ECG HRV and ECG HRV signals, suggesting a promising degree of similarity and correspondence. The findings highlight the feasibility of extracting M-ECG and computing HRV directly from raw MEG data. These insights hold the potential to enhance multimodal neuroimaging methodologies and further elucidate the intricate interplay between brain activity and cardiovascular function. The potential of HRV as a biomarker for brain disorders could improve diagnostic accuracy, prognostic assessment, and therapeutic strategies, particularly in neurological disorders with centrally mediated autonomic dysfunction.

## 1. Introduction

Magnetoencephalography (MEG) is a non-invasive neurophysiological technique that measures the magnetic fields generated by neuronal activity in the brain (Singh, 2014) and can be used to produce images of brain activity. It offers excellent temporal resolution, allowing for the observation of the complex dynamics of the brain on a submillisecond scale (Singh, 2014). MEG is used to detect brain oscillations generated across different regions of the brain (Fred et al., 2022).

MEG scanners are typically housed in magnetically-shielded rooms for enhanced passive noise reduction (Singh, 2014; Vrba & Wilson, 2007) and incorporate reference sensors, such as magnetometers or gradiometers, positioned outside the main sensor array, which capture different aspects of the environmental magnetic field and subsequently facilitate noise cancellation techniques (Singh, 2014; Vrba & Robinson, 2001; Vrba & Wilson, 2007). While these reference sensors are primarily used to monitor and compensate for ambient magnetic noise, they can also detect cardiac and other muscular activities (Hari & Salmelin, 2012; Jousmäki & Hari, 1996; Proudfoot et al., 2014; Sander et al., 2002).

MEG analyses usually entail the preprocessing of data to remove artifacts from sources such as heartbeat, eye movements, and sources of environmental noise (Breuer et al., 2014; Escudero et al., 2007) to improve the accuracy of neuroimaging results and to increase the signal-to-noise ratio (Gonzalez-Moreno et al., 2014; Treacher et al., 2021). There are various computational methods to remove cardiac artifacts from MEG data, including independent component analysis (ICA), signal filtering methods such as high-pass, low-pass, or bandpass filtering, signal space separation (SSS), signal space projection (SSP), beamforming, machine learning, or a combination thereof (Escudero et al., 2007; Hasasneh et al., 2018; Körber et al., 2001; Litvak et al., 2010; Taulu et al., 2004; Treacher et al., 2021; van Driel et al., 2019; Vrba & Robinson, 1999). Among the methods listed, ICA is commonly used for removing cardiac artifacts from MEG data. ICA is popular because it can effectively separate sources of neural activity from artifacts, including cardiac signals, based on their statistical independence. ICA does not require prior knowledge of the artifact waveform or timing, making it a versatile and widely applicable technique in MEG data analysis (Escudero et al., 2007).

The ability to remove these artifacts also raises the possibility that useful information may be obtained by isolating them. In particular, heart rate variability (HRV) serves as a significant biomarker in the study of brain disorders (Arakaki et al., 2023). The autonomic nervous system (ANS), which regulates heart rate and other physiological functions, is closely linked to brain function, and dysfunction of the ANS is associated with various neurological and psychiatric disorders, including epilepsy, Parkinson's disease, and mood disorders (Arakaki et al., 2023; Arnao et al., 2020; Bassett, 2015; Myers et al., 2018). Therefore, changes in HRV can provide insights into ANS activity and its dysregulation in brain disorders.

Many brain disorders, such as stroke, dementia, traumatic brain injury (TBI), post-traumatic stress disorder (PTSD), and sudden unexpected death in epilepsy (SUDEP) are associated with an increased risk of cardiovascular complications (Baysal-Kirac et al., 2017; Deckers et al., 2017; Samuels, 2007; Schnabel et al., 2021; Song et al., 2021; Stewart et al., 2022; Tan et al., 2011; Wolters et al., 2018). Monitoring HRV could serve as a diagnostic marker for specific brain disorders such as PTSD; it could also predict disease progression and outcomes (de Faria Cardoso et al., 2022; Liddell et al., 2016; Turcu et al., 2023).

The RR interval, derived from the time between consecutive R waves in an electrocardiogram (ECG), serves as a crucial measure for assessing HRV and evaluating the rhythm and regularity of the heartbeat (Lanfranchi & Somers, 2011; Peltola, 2012). In addition, in HRV analysis, the very-low-frequency (VLF), low-frequency (LF), and high-frequency (HF) bands represent specific frequency ranges within the HRV spectrum (Usui & Nishida, 2017). The VLF band typically spans frequencies from 0.0033 to 0.04 Hz, reflecting the activity of the central nervous system (CNS) (Armstrong et al., 2022; McCraty, 2022). The LF band typically spans frequencies from 0.04 to 0.15 Hz, reflecting sympathetic and parasympathetic modulation (Batchinsky et al., 2007; Cabiddu et al., 2012; Duprey et al., 2019; Scavone et al., 2021), while the HF band covers frequencies from 0.15 to 0.4 Hz, primarily indicating parasympathetic (vagal) activity and respiratory sinus arrhythmia (Batchinsky et al., 2007; Draghici & Taylor, 2016; Duprey et al., 2019; Hayano & Yuda, 2021; Scavone et al., 2021). These bands are utilized to evaluate the balance between sympathetic and parasympathetic nervous system activity. This suggests that HRV monitoring may be a useful tool to assess the effects of treatments for brain disorders on cardiovascular function, enhancing understanding of the complex interplay between the brain and the cardiovascular system and holding promise for improving diagnosis, prognosis, and therapeutic strategies.

Considering the importance of the ECG signal and its potential as a biomarker for a range of brain disorders, accurately extracting the ECG signal from raw MEG data and subsequently deriving HRV from it can offer valuable insights (Godwin et al., 2024). While the implementation of the ICA method alone can help with this goal (Godwin et al., 2024), in this study, a novel algorithm is proposed, which is designed to detect and extract the ECG signal, as well as compute HRV, from MEG data accurately and reliably using either the ICA method or the MEG reference sensors. This algorithm is used to detect valid R peaks, automatically identify outlier R peaks, and correct outlier RR intervals to compute HRV reliably. The performance of this algorithm is evaluated, and the quality of the extracted ECG signals from MEG, along with their computed HRV measures, are compared with an actual recorded ECG signal. Signal processing methods in the time, frequency, and time-frequency domains along with statistical tests such as Spearman’s correlation, root mean square error, mean absolute error, Bland-Altman mean difference, and Mann-Whitney U Test are employed to assess the similarities across the signals. The results underscore the feasibility of utilizing raw MEG data for ECG and HRV extraction.

## 2. Materials and Methods

### 2.1. Data

This study used Brainstorm’s MEG auditory dataset, specifically the CTF dataset (Bock et al., 2013), with prior consent from Brainstorm for research use of this publicly available dataset. The dataset comprises data from a single participant recorded using the CTF 275-channel MEG system equipped with 26 reference sensors and 274 MEG axial gradiometers (one channel was disabled). Two acquisition runs of 6 minutes each were conducted with a 2400 Hz sampling rate and an anti-aliasing low-pass filter set at 600 Hz. The data files were saved with the 3^rd^-order gradient correction applied offline. During the acquisitions, the participant was stimulated binaurally using intra-aural earphones equipped with air tubes plus transducers. Simultaneously, the participant's ECG data were recorded using a separate bipolar channel.

Additionally, the Open MEG Archive (OMEGA) resting-state sample dataset (Niso et al., 2016) was used in this study, which includes recordings from five participants using a CTF 275-channel MEG system with 26 reference sensors and 270 MEG axial gradiometers. One acquisition run of 5 minutes length was conducted with a 2400 Hz sampling rate and an anti-aliasing low-pass filter set at 600 Hz. The data files were saved with or without 3^rd^-order gradient compensation. During the acquisition, participants rested with their eyes open, and ECG data were recorded using a separate bipolar channel.

The study received approval from the Atrium Health Wake Forest Baptist institutional review board (IRB00110720). These datasets enabled an assessment of the accuracy of the extracted ECG and HRV from the MEG data in comparison to the simultaneously recorded ECG signal.

### 2.2. Data Preprocessing

MNE-Python, an open-source Python package tailored for exploring, visualizing, and analyzing various types of human neurophysiological data such as MEG, EEG, etc., was utilized to preprocess the raw MEG data (Gramfort et al., 2013; Larson et al., 2024). Other libraries used extensively in all stages of data analysis include Matplotlib, NumPy, and SciPy (Harris et al., 2020; Hunter, 2007).

During preprocessing (Fig. 1), the recorded ECG signal was extracted and saved as a separate channel. To ensure clarity, the separately recorded ECG signal is referred to as the independent ECG (I-ECG), and the ECG signal extracted from the MEG data as MEG ECG (M-ECG). These abbreviations are used throughout to maintain clarity and coherence. To clarify, while the M-ECG signal originates from the raw MEG signals, it should not be confused with magnetocardiography (MCG), which is a distinct technique used to measure the magnetic fields produced by the electrical activity of the heart. MCG specifically focuses on cardiac activity to assess heart function, detect abnormalities, and study cardiac disorders (Fenici et al., 2005).

**Fig. 1.**
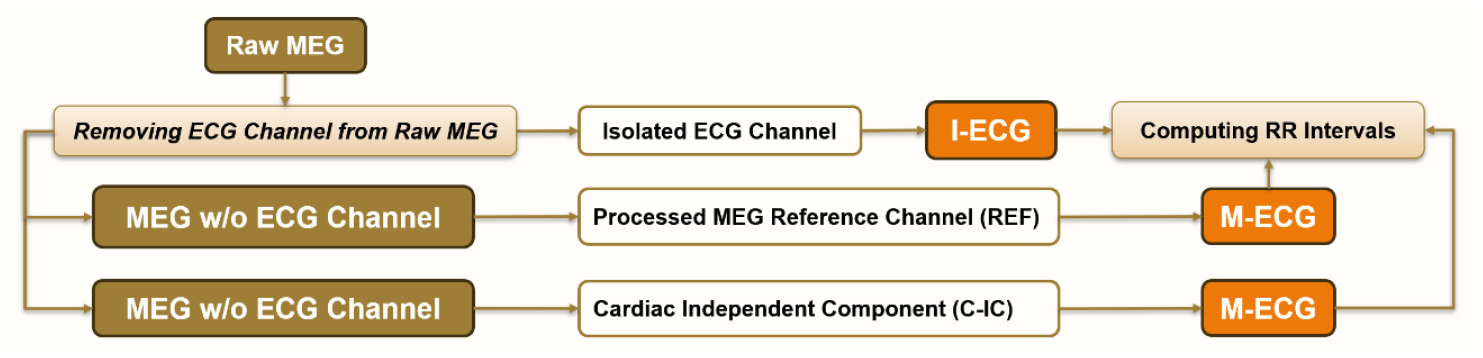
Preprocessing pipeline.

**Fig. 2.**
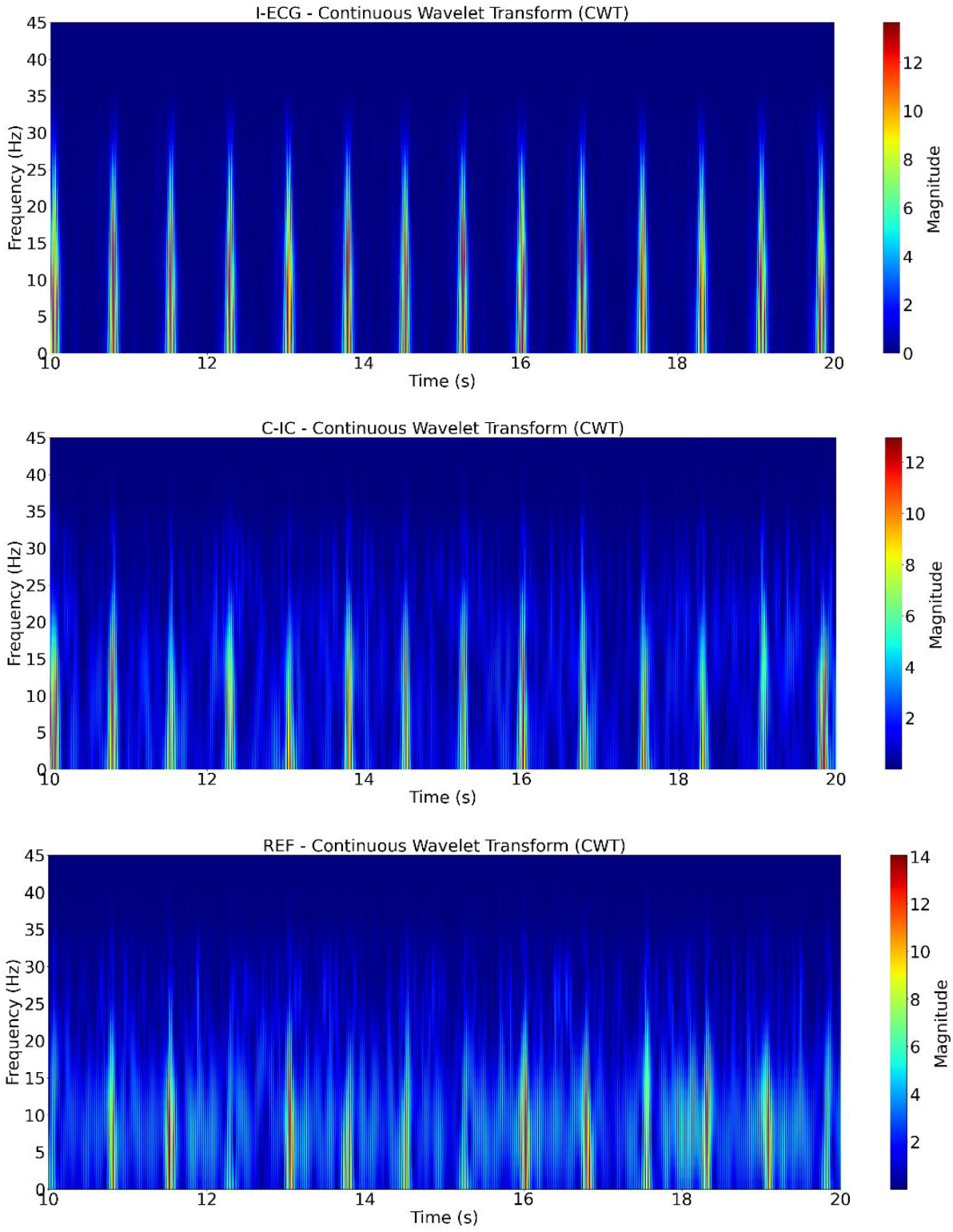
CWT analysis of M-ECG and I-ECG Signals using the Morlet wavelet (participant 1).

Three participants from the OMEGA resting-state sample dataset were excluded from the study due to high levels of artifact and noise in the MEG data, which compromised the quality of the recordings, while two participants were retained. Along with one participant from the Brainstorm dataset, a total of three participants were included in the study. The raw MEG data were bandpass filtered from 0.5 - 45 Hz to retain relevant ECG frequency information (Wasimuddin et al., 2020). A total of 27 channels from the right and left temporal regions (MRT/MLT 31-58), exhibiting a strong cardiac signature, were selected. Through an iterative process, the ICA components of 15 proved sufficient in detecting the cardiac component. The component showing the most robust cardiac activity was then designated as a virtual channel, referred to as the cardiac independent component (C-IC) (Fig. 1). Similarly, the reference channel displaying the most robust cardiac activity was selected. To ensure clarity, the selected reference channel is referred to as REF (Fig. 1). Together, C-IC and REF constitute the M-ECG channels.

All three channels, I-ECG, C-IC, and REF were baseline corrected and normalized. C-IC and REF underwent discrete wavelet transform (DWT), using the PyWavelets library (Lee et al., 2019), to decompose the signals into wavelet coefficients. The coefficients from level 0, capturing the broadest, low-frequency features associated with the cardiac signal, were retained for reconstruction, emphasizing the cardiac-related components.

### 2.3. Computing RR Intervals

To automatically detect valid R-peaks in the I-ECG and M-ECG signals, a tailored algorithm was employed. Initially, invalid peaks were filtered out using a predefined low and high threshold (e.g., σ > 2). This threshold could vary depending on data quality and noise levels. The process ensured the retention of consistent, reliable R-peaks while minimizing noise and artifacts (Fig. 3, Supplemental Figs. 1-2). Subsequently, to address potential distortions in RR interval computation due to outlier removal, outliers were identified by calculating thresholds based on the mean and standard deviation of the RR intervals, specifically targeting deviations beyond a predefined low and high threshold (e.g., σ > 2.5). These outliers were set to zero to mitigate their influence. The continuity of the RR interval signal was restored through linear interpolation, filling gaps with estimated values based on surrounding data points (Fig. 4). This approach ensured a smooth, continuous RR interval series, facilitating more accurate analysis while preserving the integrity of the underlying signal. This two-step process—detection and correction—provided a robust method for RR interval computation, particularly in the presence of artifacts and outliers, enhancing the reliability of HRV analyses.

**Fig. 3.**
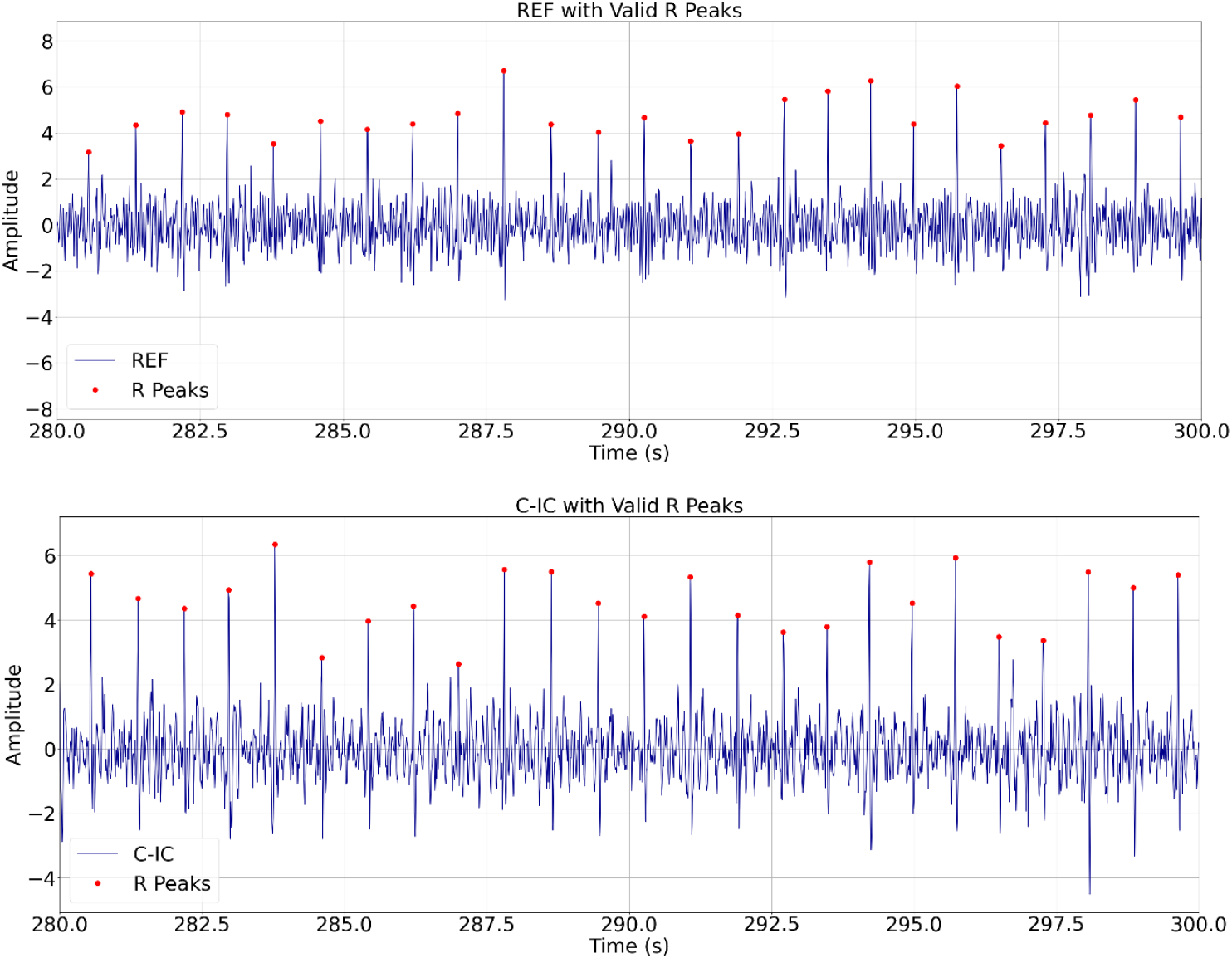
Automatically detected valid R peaks for REF and C-IC for participant 1 (280 – 300 s). Red dots indicate the detected valid peaks on each signal.

**Fig. 4.**
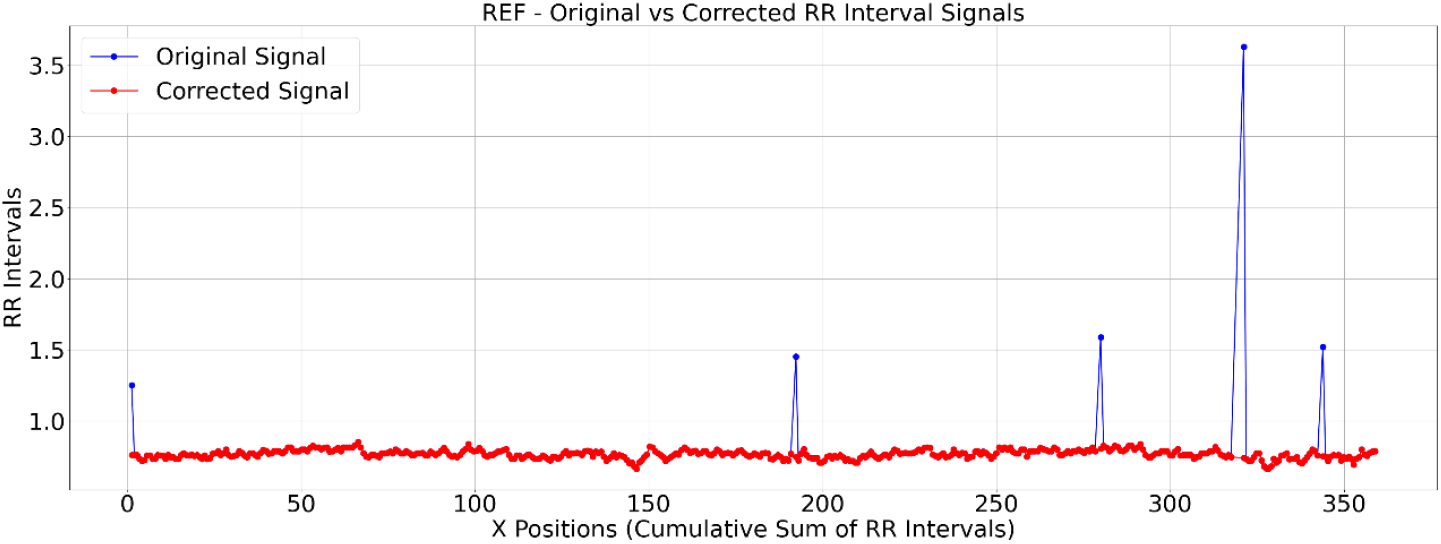
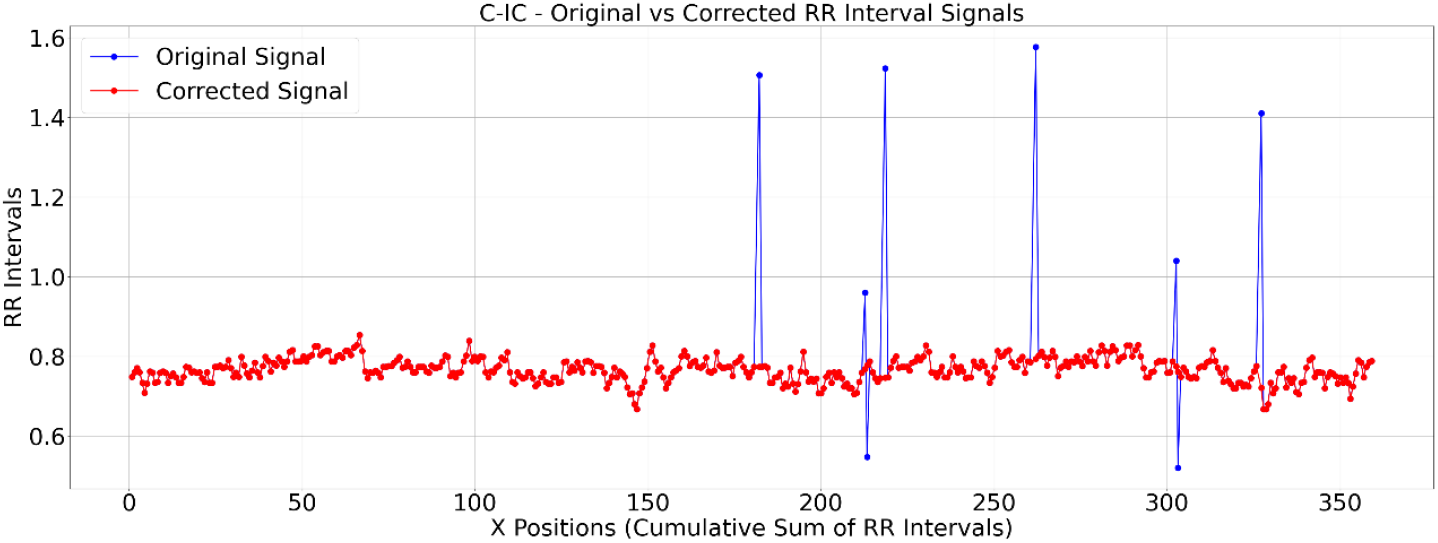
Original vs corrected RR intervals for REF and C-IC for participant 1 (0 – 360 s). The corrected signal is shown in red, while the uncorrected signal appears in blue.

### 2.4. Time Domain

To allow for clearer visualization of consistent peak patterns between the M-ECG and I-ECG signals, peak-to-peak signal averaging around R-peaks was computed for I-ECG and M-ECG signals (Fig. 5, Supplemental Figs. 3-4). Additionally, to compare time series sequences between I-ECG RR intervals and M-ECG RR intervals, dynamic time warping (DTW) was computed (Fig. 6, Supplemental Figs. 5-6) using the DTAIDistance Python library (Meert et al., 2022). DTW is a powerful technique that allows for comparing time series data that may vary in length or timing by finding an optimal alignment path between the sequences, enabling comparison and similarity measurement even when the sequences have different lengths or rates of change (Senin, 2008).

**Fig. 5.**
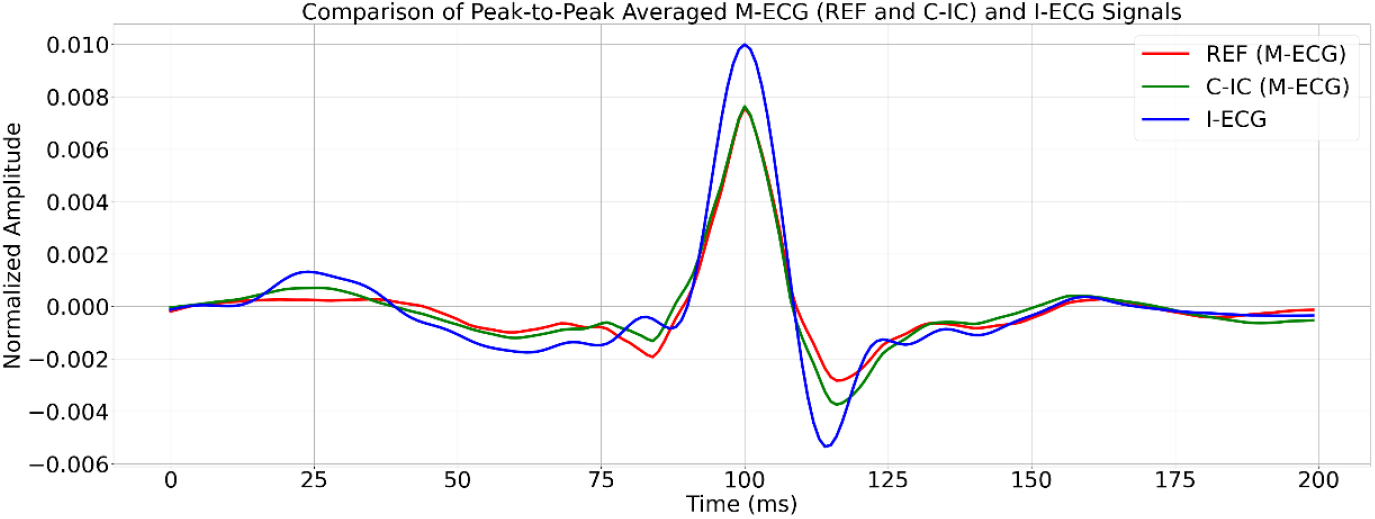
Comparison of averaged peak-to-peak REF, C-IC, and I-ECG signals for participant 1. REF is shown in red, C-IC in green, and I-ECG in blue.

**Fig. 6.**
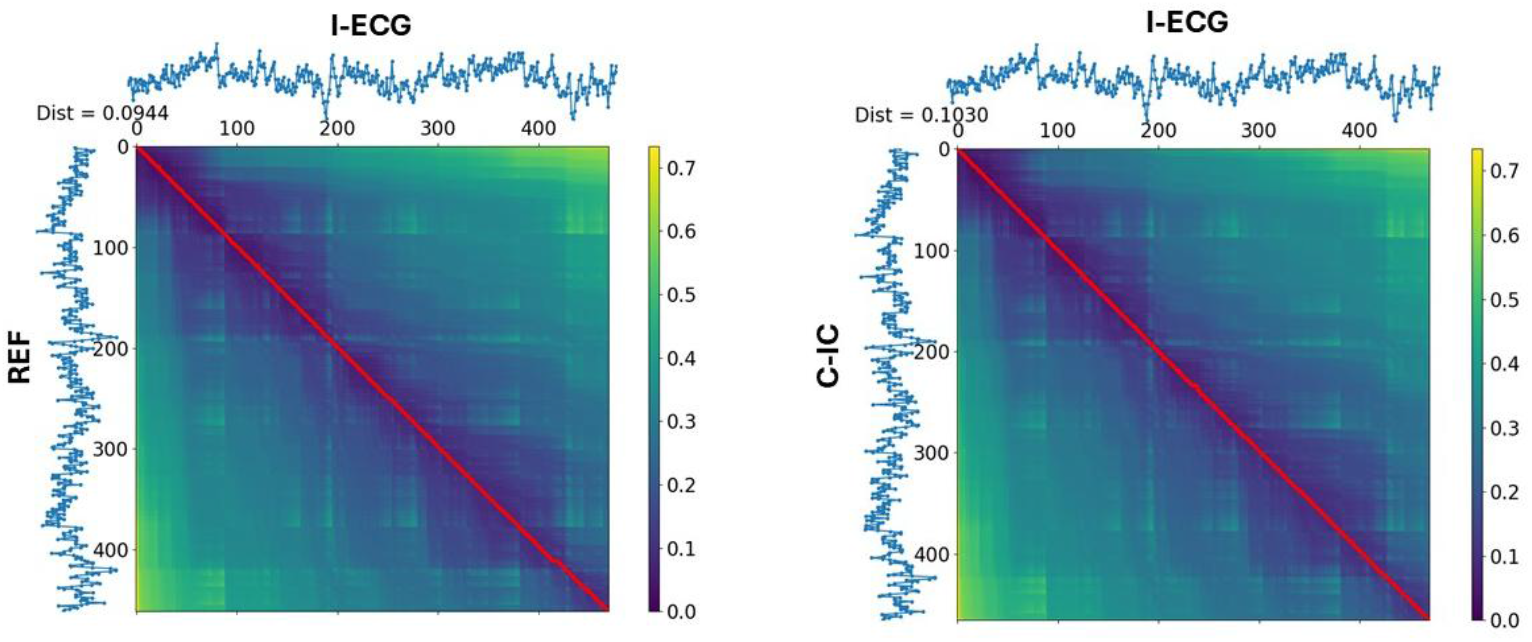
DTW of RR intervals with I-ECG vs REF (left) and I-ECG vs C-IC (right) for participant 1. The lower the DTW value, the more similar the two signals are, with a value closer to zero indicating a better alignment and higher similarity.

Widely-used time-domain HRV indices, including the standard deviation of normal-to-normal intervals (SDNN), which reflects overall variability, and the root mean square of successive differences (RMSSD), which estimates vagally mediated changes in HRV (Munoz et al., 2015; Shaffer & Ginsberg, 2017), were computed for both I-ECG and M-ECG signals (Table 1). Comparing SDNN and RMSSD between the two signals allowed for assessing their similarity in capturing HRV.

**Table 1.**
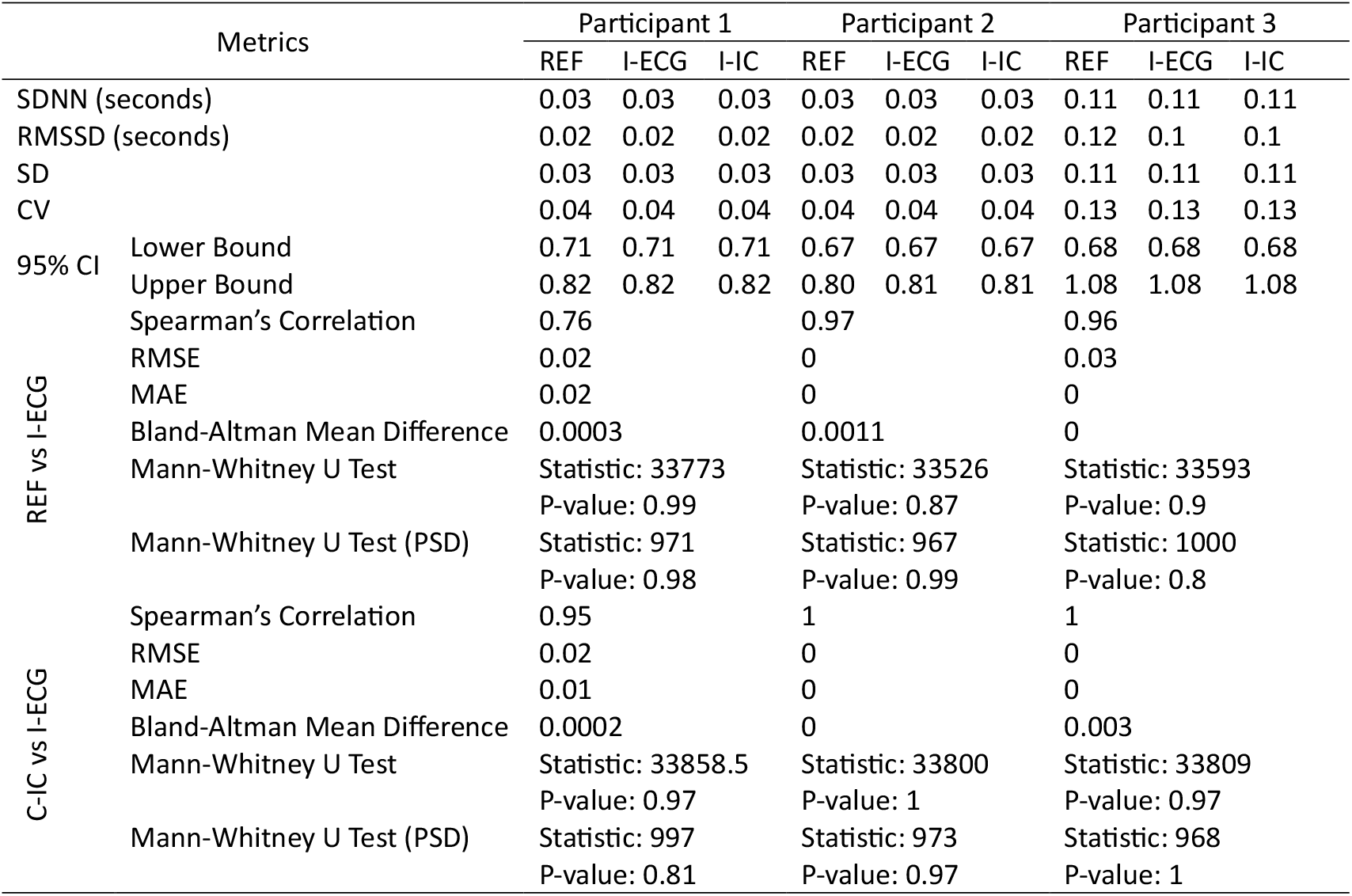
Detailed statistical comparison of RR intervals derived from M-ECG signals (REF and C-IC) against those from I-ECG for all 3 participants (0 – 200 s).

### 2.5. Frequency and Time-Frequency Domains

To investigate transient events, frequency modulation, and other dynamic characteristics of I-ECG and M-ECG signals, the continuous wavelet transform (CWT) was employed due to its high time-frequency resolution. This method, implemented using the Scipy and PyWavelet libraries (Lee et al., 2019; Virtanen et al., 2020), allowed for the examination of signals with localized features that change in frequency over time. To visualize the results of these analyses, spectrograms of time and frequency were generated (Fig. 2).

Fig. 2 presents the CWT analysis of I-ECG and M-ECG signals using the Morlet wavelet for participant 1. The plots depict the time-frequency decomposition of the signals over a 20-second duration. Recurrent high-power bursts in the plots likely correspond to specific R waves, reflecting cardiac activity.

Since frequency and time-frequency analyses often assume evenly sampled data, the uneven sampling of RR intervals, due to natural HRV, posed a potential limitation (Hashimoto et al., 2013). To mitigate this, Cubic Spline interpolation was used to convert the unevenly sampled RR intervals into evenly sampled signals, facilitating accurate frequency and time-frequency analyses. Cubic Spline interpolation achieves this by fitting a piecewise cubic polynomial between adjacent data points (de Boor, 2001; Dyer & Dyer, 2001), ensuring smoothness and continuity in the interpolated signal.

To assess the frequency-domain properties and interrelations of the I-ECG and M-ECG interpolated RR intervals, power spectral density (PSD) analysis was performed to reveal the spectral distribution of each RR interval signal (Fig. 7, Supplemental Figs 7-8).

### 2.6. Statistical Analyses

To measure the similarity between I-ECG and M-ECG RR intervals, several statistical metrics were computed. Standard deviation (SD) was used to assess the variability and precision of RR interval measurements derived from M-ECG and I-ECG. To quantify the average difference between I-ECG and M-ECG RR intervals, root mean squared error (RMSE) and mean absolute error (MAE) were calculated, while the coefficient of variation (CV) was calculated to measure the relative variability of both sets of intervals. 95% confidence intervals (CI) were computed to estimate the range within which the true RR interval parameters likely fall, assessing the precision and reliability of the metrics. The Spearman rank correlation coefficient was also computed to gauge the correlation strength of M-ECG and I-ECG RR intervals.

To determine if RR intervals from I-ECG and M-ECG were drawn from populations with the same distribution, the Mann-Whitney U test was performed. This test assessed whether there were significant differences (temporally and frequency-wise) in the distributions of RR intervals between I-ECG and M-ECG (Table 1). Statistical analyses were performed using Scipy and Scikit-learn libraries (Pedregosa et al., 2011; Virtanen et al., 2020). Finally, Bland-Altman analysis (Martin Bland & Altman, 1986) was conducted to compare the mean differences between sets of RR intervals derived from M-ECG and I-ECG, identifying any systematic bias or proportional differences between the two measurement methods (Table 1). The custom analytical script is made available on GitHub (https://github.com/izadysadr/MEG-HRV-Extraction) to facilitate replication.

## 3. Results

The low/high thresholding approach effectively filtered out invalid peaks, retaining only consistent and physiologically plausible R peaks while excluding artifacts (Fig. 3, Supplemental Figs. 1-2). Additionally, the algorithm identified and interpolated outliers in the RR intervals (Fig. 4), ensuring a smooth and continuous time series. Together, these steps helped mitigate the impact of artifacts and outliers, thereby enhancing the reliability of the computed RR interval sequences.

A comparison of peak-to-peak signal averaging between I-ECG and M-ECG revealed that while the peak amplitude of I-ECG is slightly higher than that of M-ECG, the alignment of the average peak amplitude between the signals indicates a match in capturing the R-peaks (Fig. 5, Supplemental Figs. 3-4).

The DTW analysis revealed a DTW distance of approximately 0.0944 seconds between REF and I-ECG RR intervals and 0.1310 seconds between C-IC and I-ECG RR intervals (Fig. 6). The nearly straight alignment path observed in Fig. 6 and Supplemental Figs. 5-6 indicates a close temporal alignment between M-ECG and I-ECG RR intervals.

The PSD analysis of REF, C-IC, and I-ECG across VLF, LF, and HF bands (Fig. 7, Supplemental Figs 7-8) demonstrates that REF and C-IC RR intervals retain the spectral distributions observed in the I-ECG RR intervals across these frequency bands.

The statistical comparison of the RR intervals derived from M-ECG (REF and C-IC) against those from I-ECG for all 3 participants (Table 1) highlights the similarity and strong agreement between the two sets of signals. The SDNN, RMSSD, SD, CV, and CI values for both REF and C-IC are nearly identical to those for I-ECG in each participant. Both REF and C-IC exhibit high Spearman’s rank correlation values with I-ECG across participants, reflecting strong linear relationships. In addition, RMSE and MAE values appear close to zero in each participant, confirming the similarity between the M-ECG and I-ECG RR intervals.

The Bland-Altman analysis revealed negligible mean differences between the REF and I-ECG, as well as between the C-IC and I-ECG for each participant, suggesting high agreement between the signals. Furthermore, the Mann-Whitney U tests yielded no significant statistical differences between the signal sets in each participant.

**Fig. 7.**
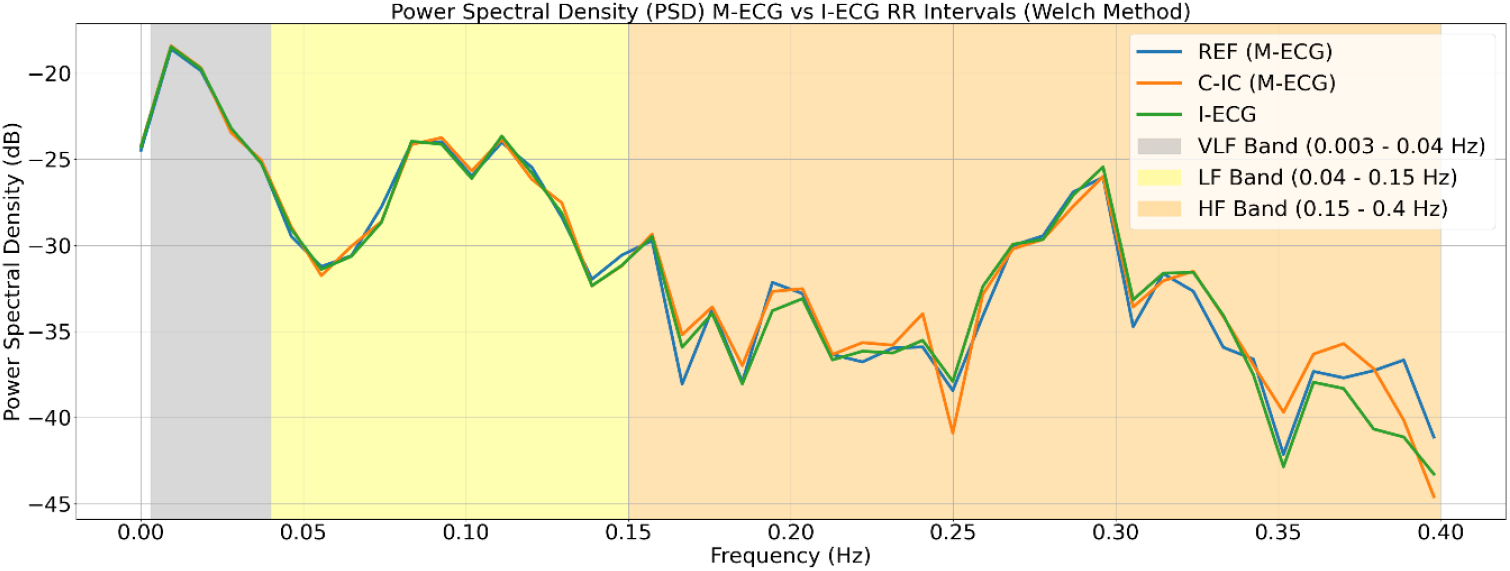
PSD between REF, C-IC, and I-ECG RR intervals for participant 1. VLF, LF, and HF bands are analyzed across REF (blue), C-IC (orange), and I-ECG (green) signals.

## 4. Discussion

This study aimed to evaluate the efficacy of M-ECG in capturing cardiac activity and to compare these signals to the I-ECG recording. Using a rigorous preprocessing and analysis pipeline, multiple aspects of the signals, including RR interval computation, time-domain indices, frequency-domain characteristics, and statistical testing were examined. The results highlight the potential of M-ECG signals to serve as a reliable surrogate for I-ECG in specific applications, such as HRV analysis and cardiovascular research.

The preprocessing pipeline employed for extracting the cardiac signals from MEG data was effective in isolating cardiac components. The study successfully identified and designated the cardiac signals using REF and C-IC as M-ECG channels. The use of bandpass filtering of 0.5–45 Hz, baseline correction, normalization, and application of DWT ensured that only the most relevant cardiac-related features were retained. This approach allowed for robust reconstruction of cardiac-related components in REF and C-IC channels while suppressing noise.

The tailored algorithm for detecting R-peaks demonstrated high precision in excluding invalid peaks and minimizing artifacts. The dynamic thresholding approach ensured that physiologically plausible peaks were retained, while outlier removal and linear interpolation provided a smooth and continuous RR interval series. These steps allowed for maintaining the integrity of HRV analyses, particularly when working with noisy or artifact-prone data.

The DTW analysis provided insights into the temporal alignment of RR intervals between M-ECG and I-ECG signals, reflected by nearly straight alignment paths and low DTW distances for both REF and C-IC when compared to I-ECG, highlighting the temporal similarity between these signals.

The time-domain HRV indices, including SDNN and RMSSD, were nearly identical between M-ECG and I-ECG signals, demonstrating that M-ECG signals can capture overall HRV and vagally-mediated HRV changes with high fidelity. In addition, peak-to-peak averaging demonstrated similarity between M-ECG and I-ECG signals, underscoring the capability of M-ECG signals to faithfully replicate cardiac dynamics in I-ECG.

The PSD analysis revealed that the M-ECG-derived RR intervals accurately reflect the spectral distribution of I-ECG RR intervals across VLF, LF, and HF bands. This similarity underscores the ability of M-ECG signals to capture the autonomic balance and other physiological processes reflected in these frequency bands. The use of CWT further allowed for high-resolution examination of transient events and dynamic characteristics. The Morlet wavelet-based CWT analysis showed time-frequency patterns corresponding to cardiac activity.

The statistical analyses provided further evidence for the equivalence of M-ECG and I-ECG RR intervals. The Bland-Altman analysis revealed minimal mean differences between the two methods, indicating negligible systematic bias. The high Spearman’s rank correlation and low RMSE and MAE values further supported the agreement between the signal sets. Additionally, the Mann-Whitney U test in the time and frequency domains confirmed that the RR intervals and their PSD values from M-ECG and I-ECG were not significantly different, suggesting that M-ECG and I-ECG RR intervals are highly similar in both time-domain and frequency-domain characteristics.

## 5. Limitations and Conclusion

The efficacy of the C-IC method is contingent upon the availability of sufficient MEG data for identifying cardiac activity within the dataset. For this study, MEG data with a duration of 360 seconds were used, and shorter durations of MEG recording might not encompass enough cardiac activity for accurate detection through the ICA method. Through an iterative process, it was determined that the ICA method requires a minimum of 6 seconds of recorded MEG data to identify cardiac activity correctly, as evidenced by the corresponding topoplot of the cardiac component. Additionally, if the MEG data are excessively noisy, the resulting REF and C-IC signals may be compromised, potentially leading to less reliable results. Finally, the small sample size in this study represents a limitation, and future research should involve larger cohorts to further validate the robustness of the results.

The M-ECG extraction method can be adapted for use with various MEG systems, allowing for the extraction of normal and abnormal HRV, which can serve as a valuable biomarker for neurological disorders closely linked to ANS dysregulation. Extracting M-ECG facilitates the retrospective analysis of the relationship between HRV and neuronal function in patients with cardiac and neurological conditions, providing deeper insights into disease mechanisms and advancing both diagnostic and therapeutic strategies.

In conclusion, this study introduces a novel algorithm for extracting cardiac activity from MEG data to accurately approximate HRV measures from actual ECG recordings. Through analysis across various domains, including time, frequency, and time-frequency, as well as statistical assessments, the robustness and comparability of both C-IC and REF in capturing HRV and cardiac dynamics was demonstrated. The findings suggest the potential clinical utility of M-ECG in exploring brain-heart interactions and advancing neurocardiac research.

## Acknowledgments

This study was supported by grant AA016852 to Dwayne W. Godwin. Aqil Izadysadr, Cormac A. O`Donovan, Gautam Popli, Hamideh Sadat Bagherzadeh, and Jennifer R. Stapleton-Kotloski were supported by the Department of Neurology.

## CRediT authorship contribution statement

**Aqil Izadysadr:** Formal analysis, Methodology, Software, Validation, Data curation, Visualization, Writing – original draft, Writing – review and editing. **Hamideh Sadat Bagherzadeh:** Formal analysis, Writing – original draft, Writing – review and editing, Project administration. **Jennifer R. Stapleton-Kotloski:** Supervision, Methodology, Validation, Writing – review and editing. **Gautam Popli:** Writing – review and editing. **Cormac A. O'Donovan:** Writing – review and editing. **Dwayne W. Godwin:** Conceptualization, Validation, Supervision, Methodology, Funding acquisition, Resources, Project administration, Writing – review and editing

## Appendix A. Supplementary Data

The supplementary data for this article are as follows: (Attached Document)

## References

Arakaki, X., Arechavala, R. J., Choy, E. H., Bautista, J., Bliss, B., Molloy, C., Wu, D. A., Shimojo, S., Jiang, Y., Kleinman, M. T., & Kloner, R. A. (2023). The connection between heart rate variability (HRV), neurological health, and cognition: A literature review. Frontiers in Neuroscience, 17, 1055445. 10.3389/FNINS.2023.1055445

Armstrong, R., Wheen, P., Brandon, L., Maree, A., & Kenny, R. A. (2022). Heart rate: control mechanisms, pathophysiology and assessment of the neurocardiac system in health and disease. QJM: An International Journal of Medicine, 115(12), 806–812. 10.1093/QJMED/HCAB016

Arnao, V., Cinturino, A., Mastrilli, S., Buttà, C., Maida, C., Tuttolomondo, A., Aridon, P., & D’Amelio, M. (2020). Impaired circadian heart rate variability in Parkinson’s disease: A time-domain analysis in ambulatory setting. BMC Neurology, 20(1), 1–5. 10.1186/S12883-020-01722-3/FIGURES/2

Bassett, D. (2015). A literature review of heart rate variability in depressive and bipolar disorders. http://dx.doi.org.wake.idm.oclc.org/10.1177/0004867415622689, 50(6), 511–519. 10.1177/0004867415622689

Batchinsky, A. I., Cooke, W. H., Kuusela, T. A., Jordan, B. S., Wang, J. J., & Cancio, L. C. (2007). Sympathetic nerve activity and heart rate variability during severe hemorrhagic shock in sheep. Autonomic Neuroscience, 136(1–2), 43–51. 10.1016/J.AUTNEU.2007.03.004

Baysal-Kirac, L., Serbest, N. G., Şahin, E., Dede, H. Ö., Gürses, C., Gökyiğit, A., Bebek, N., Bilge, A. K., & Baykan, B. (2017). Analysis of heart rate variability and risk factors for SUDEP in patients with drug-resistant epilepsy. Epilepsy & Behavior : E&B, 71(Pt A), 60–64. 10.1016/J.YEBEH.2017.04.018

Bock, E., Donhauser, P., Tadel, F., & Baillet, S. (2013). Dataset Auditory - Brainstorm. Montreal Neurological Institute.

Breuer, L., Dammers, J., Roberts, T. P. L., & Shah, N. J. (2014). Ocular and cardiac artifact rejection for real-time analysis in MEG. Journal of Neuroscience Methods, 233, 105–114. 10.1016/J.JNEUMETH.2014.06.016

Cabiddu, R., Cerutti, S., Viardot, G., Werner, S., & Bianchi, A. M. (2012). Modulation of the sympatho-vagal balance during sleep: Frequency domain study of heart rate variability and respiration. Frontiers in Physiology, 3 MAR, 20851. 10.3389/FPHYS.2012.00045/BIBTEX

de Boor, C. (2001). Spline Interpolation. 171–205. 10.1007/978-1-4612-6333-3_13

de Faria Cardoso, C., Ohe, N. T., Bader, Y., Afify, N., Al-Homedi, Z., Alwedami, S. M., O’Sullivan, S., Campos, L. A., & Baltatu, O. C. (2022). Heart Rate Variability Indices as Possible Biomarkers for the Severity of Post-traumatic Stress Disorder Following Pregnancy Loss. Frontiers in Psychiatry, 12. 10.3389/FPSYT.2021.700920/FULL

Deckers, K., Schievink, S. H. J., Rodriquez, M. M. F., Van Oostenbrugge, R. J., Van Boxtel, M. P. J., Verhey, F. R. J., & Kö Hler, S. (2017). Coronary heart disease and risk for cognitive impairment or dementia: Systematic review and meta-analysis. Journals.Plos.OrgK Deckers, SHJ Schievink, MMF Rodriquez, RJ van Oostenbrugge, MPJ van BoxtelPloS One, 2017.journals.Plos.Org, 12(9). 10.1371/journal.pone.0184244

Draghici, A. E., & Taylor, J. A. (2016). The physiological basis and measurement of heart rate variability in humans. Journal of Physiological Anthropology, 35(1), 1–8. 10.1186/S40101-016-0113-7/FIGURES/3

Duprey, E. B., Oshri, A., & Liu, S. (2019). Childhood Maltreatment, Self-esteem, and Suicidal Ideation in a Low-SES Emerging Adult Sample: The Moderating Role of Heart Rate Variability. Archives of Suicide Research, 23(2), 333–352. 10.1080/13811118.2018.1430640

Dyer, S. A., & Dyer, J. S. (2001). Cubic-Spline Interpolation: Part 1. IEEE Instrumentation and Measurement Magazine, 4(1), 44–46. 10.1109/5289.911175

Escudero, J., Hornero, R., Abasolo, D., Fernandez, A., & Lopez-Coronado, M. (2007). Artifact removal in magnetoencephalogram background activity with independent component analysis. IEEE Transactions on Biomedical Engineering, 54(11), 1965–1973. 10.1109/TBME.2007.894968

Fenici, R., Brisinda, D., & Meloni, A. M. (2005). Clinical application of magnetocardiography. Expert Review of Molecular Diagnostics, 5(3), 291–313. 10.1586/14737159.5.3.291

Fred, A. L., Kumar, S. N., Haridhas, A. K., Ghosh, S., Bhuvana, H. P., Sim, W. K. J., Vimalan, V., Givo, F. A. S., Jousmäki, V., Padmanabhan, P., & Gulyás, B. (2022). A Brief Introduction to Magnetoencephalography (MEG) and Its Clinical Applications. Brain Sciences 2022, Vol. 12, Page 788, 12(6), 788. 10.3390/BRAINSCI12060788

Godwin, R. C., Flood, W. C., Hudson, J. P., Benayoun, M. D., Zapadka, M. E., Melvin, R. L., & Whitlow, C. T. (2024). Automated extraction of heart rate variability from magnetoencephalography signals. Heliyon, 10, e26664. 10.1016/j.heliyon.2024.e26664

Gonzalez-Moreno, A., Aurtenetxe, S., Lopez-Garcia, M. E., del Pozo, F., Maestu, F., & Nevado, A. (2014). Signal-to-noise ratio of the MEG signal after preprocessing. Journal of Neuroscience Methods, 222, 56–61. 10.1016/J.JNEUMETH.2013.10.019

Gramfort, A., Luessi, M., Larson, E., Engemann, D. A., Strohmeier, D., Brodbeck, C., Goj, R., Jas, M., Brooks, T., Parkkonen, L., & Hämäläinen, M. (2013). MEG and EEG data analysis with MNE-Python. Frontiers in Neuroscience, 7(7 DEC), 70133. 10.3389/FNINS.2013.00267/BIBTEX

Hari, R., & Salmelin, R. (2012). Magnetoencephalography: From SQUIDs to neuroscience. Neuroimage 20th anniversary special edition. NeuroImage, 61(2), 386–396. 10.1016/J.NEUROIMAGE.2011.11.074

Harris, C. R., Millman, K. J., van der Walt, S. J., Gommers, R., Virtanen, P., Cournapeau, D., Wieser, E., Taylor, J., Berg, S., Smith, N. J., Kern, R., Picus, M., Hoyer, S., van Kerkwijk, M. H., Brett, M., Haldane, A., del Río, J. F., Wiebe, M., Peterson, P., … Oliphant, T. E. (2020). Array programming with NumPy. Nature 2020 585:7825, 585(7825), 357–362. 10.1038/s41586-020-2649-2

Hasasneh, A., Kampel, N., Sripad, P., Shah, N. J., & Dammers, J. (2018). Deep Learning Approach for Automatic Classification of Ocular and Cardiac Artifacts in MEG Data. Journal of Engineering (United Kingdom), 2018. 10.1155/2018/1350692

Hashimoto, H., Fujiwara, K., Suzuki, Y., Miyajima, M., Yamakawa, T., Kano, M., Maehara, T., Ohta, K., Sasano, T., Matsuura, M., & Matsushima, E. (2013). Heart rate variability features for epilepsy seizure prediction. 2013 Asia-Pacific Signal and Information Processing Association Annual Summit and Conference, APSIPA 2013. 10.1109/APSIPA.2013.6694240

Hayano, J., & Yuda, E. (2021). Assessment of autonomic function by long-term heart rate variability: beyond the classical framework of LF and HF measurements. In Journal of Physiological Anthropology (Vol. 40, Issue 1, pp. 1–15). BioMed Central Ltd. 10.1186/s40101-021-00272-y

Hunter, J. D. (2007). Matplotlib: A 2D graphics environment. Computing in Science \& Engineering, 9(3), 90–95. 10.1109/MCSE.2007.55

Jousmäki, V., & Hari, R. (1996). Cardiac artifacts in magnetoencephalogram. Journal of Clinical Neurophysiology, 13(2), 172–176. 10.1097/00004691-199603000-00008

Körber, R., Storm, J.-H., Seton, H., -, al, Cai, C., Kang, H., Kirsch, H. E., Nolte, G., & Hämäläinen, M. S. (2001). Partial signal space projection for artefact removal in MEG measurements: a theoretical analysis. Physics in Medicine & Biology, 46(11), 2873. 10.1088/0031-9155/46/11/308

Lanfranchi, P. A., & Somers, V. K. (2011). Cardiovascular Physiology: Autonomic Control in Health and in Sleep Disorders. Principles and Practice of Sleep Medicine: Fifth Edition, 226–236. 10.1016/B978-1-4160-6645-3.00020-7

Larson, E., Gramfort, A., Engemann, D. A., Leppakangas, J., Brodbeck, C., Jas, M., Brooks, T., Sassenhagen, J., McCloy, D., Luessi, M., King, J.-R., Höchenberger, R., Goj, R., Favelier, G., Brunner, C., van Vliet, M., Wronkiewicz, M., Rockhill, A., Holdgraf, C., … luzpaz. (2024). MNE-Python. Zenodo. 10.5281/zenodo.10999175

Lee, G. R., Gommers, R., Waselewski, F., Wohlfahrt, K., & O’Leary, A. (2019). PyWavelets: A Python package for wavelet analysis. Journal of Open Source Software, 4(36), 1237. 10.21105/JOSS.01237

Liddell, B. J., Kemp, A. H., Steel, Z., Nickerson, A., Bryant, R. A., Tam, N., Tay, A. K., & Silove, D. (2016). Heart rate variability and the relationship between trauma exposure age, and psychopathology in a post-conflict setting. BMC Psychiatry, 16(1), 1–9. 10.1186/S12888-016-0850-5/FIGURES/2

Litvak, V., Eusebio, A., Jha, A., Oostenveld, R., Barnes, G. R., Penny, W. D., Zrinzo, L., Hariz, M. I., Limousin, P., Friston, K. J., & Brown, P. (2010). Optimized beamforming for simultaneous MEG and intracranial local field potential recordings in deep brain stimulation patients. Neuroimage, 50(4–3), 1578. 10.1016/J.NEUROIMAGE.2009.12.115

Martin Bland, J., & Altman, D. G. (1986). STATISTICAL METHODS FOR ASSESSING AGREEMENT BETWEEN TWO METHODS OF CLINICAL MEASUREMENT. The Lancet, 327(8476), 307–310. 10.1016/S0140-6736(86)90837-8

McCraty, R. (2022). Following the Rhythm of the Heart: HeartMath Institute’s Path to HRV Biofeedback. Applied Psychophysiology Biofeedback, 47(4), 305–316. 10.1007/S10484-022-09554-2/METRICS

Meert, W., Hendrickx, K., Van Craenendonck, T., Robberechts, P., Blockeel, H., & Davis, J. (2022).DTAIDistance. Zenodo. 10.5281/zenodo.7158824

Munoz, M. L., Van Roon, A., Riese, H., Thio, C., Oostenbroek, E., Westrik, I., De Geus, E. J. C., Gansevoort, R., Lefrandt, J., Nolte, I. M., & Snieder, H. (2015). Validity of (Ultra-)Short Recordings for Heart Rate Variability Measurements. PLOS ONE, 10(9), e0138921. 10.1371/JOURNAL.PONE.0138921

Myers, K. A., Sivathamboo, S., & Perucca, P. (2018). Heart rate variability measurement in epilepsy: How can we move from research to clinical practice? Epilepsia, 59(12), 2169–2178. 10.1111/EPI.14587

Niso, G., Rogers, C., Moreau, J. T., Chen, L. Y., Madjar, C., Das, S., Bock, E., Tadel, F., Evans, A. C., Jolicoeur, P., & Baillet, S. (2016). OMEGA: The Open MEG Archive. NeuroImage, 124, 1182–1187. 10.1016/J.NEUROIMAGE.2015.04.028

Pedregosa, F., Michel, V., Grisel OLIVIERGRISEL, O., Blondel, M., Prettenhofer, P., Weiss, R., Vanderplas, J., Cournapeau, D., Pedregosa, F., Varoquaux, G., Gramfort, A., Thirion, B., Grisel, O., Dubourg, V., Passos, A., Brucher, M., Perrot andÉdouardand, M., Duchesnay, A., & Duchesnay EDOUARDDUCHESNAY, Fré. (2011). Scikit-learn: Machine learning in Python. Jmlr.Org, 12, 2825–2830.

Peltola, M. A. (2012). Role of editing of R-R intervals in the analysis of heart rate variability. Frontiers in Physiology, 3 MAY, 15614. 10.3389/FPHYS.2012.00148/BIBTEX

Proudfoot, M., Woolrich, M. W., Nobre, A. C., & Turner, M. R. (2014). Magnetoencephalography. Practical Neurology, 14(5), 336. 10.1136/PRACTNEUROL-2013-000768

Samuels, M. A. (2007). The Brain–Heart Connection. Circulation, 116(1), 77–84. 10.1161/CIRCULATIONAHA.106.678995

Sander, T. H., Wübbeler, G., Lueschow, A., Curio, G., & Trahms, L. (2002). Cardiac Artifact Subspace Identification and Elimination in Cognitive MEG Data Using Time-Delayed Decorrelation. IEEE TRANSACTIONS ON BIOMEDICAL ENGINEERING, 49(4), 345.

Scavone, G., Baril, A. A., Montplaisir, J., Carrier, J., Desautels, A., & Zadra, A. (2021). Autonomic Modulation During Baseline and Recovery Sleep in Adult Sleepwalkers. Frontiers in Neurology, 12, 680596. 10.3389/FNEUR.2021.680596/BIBTEX

Schnabel, R. B., Hasenfuß, G., Buchmann, S., Kahl, K. G., Aeschbacher, S., Osswald, S., & Angermann, C. E. (2021). Heart and brain interactions: Pathophysiology and management of cardio-psycho-neurological disorders. Herz, 46(2), 138. 10.1007/S00059-021-05022-5

Senin, P. (2008). Dynamic Time Warping Algorithm Review. Science, 2007(December), 1–23.

Shaffer, F., & Ginsberg, J. P. (2017). An overview of heart rate variability metrics and norms. Frontiers in Public Health, 5, 258.

Singh, S. P. (2014). Magnetoencephalography: Basic principles. Annals of Indian Academy of Neurology, 17(SUPPL. 1). 10.4103/0972-2327.128676

Song, R., Pan, K.-Y., Xu, H., Qi, X., Buchman, A. S., Bennett, D. A., & Xu, W. (2021). Association of cardiovascular risk burden with risk of dementia and brain pathologies: a population-based cohort study. Wiley Online Library R Song, KY Pan, H Xu, X Qi, AS Buchman, DA Bennett, W XuAlzheimer’s & Dementia, 2021.Wiley Online Library, 17(12), 1914–1922. 10.1002/alz.12343

Stewart, I. J., Amuan, M. E., Wang, C. P., Kennedy, E., Kenney, K., Werner, J. K., Carlson, K. F., Tate, D. F., Pogoda, T. K., Dismuke-Greer, C. E., Wright, W. S., Wilde, E. A., & Pugh, M. J. (2022). Association Between Traumatic Brain Injury and Subsequent Cardiovascular Disease Among Post-9/11–Era Veterans. JAMA Neurology, 79(11), 1122–1129. 10.1001/JAMANEUROL.2022.2682

Tan, G., Dao, T. K., Farmer, L., Sutherland, R. J., & Gevirtz, R. (2011). Heart rate variability (HRV) and posttraumatic stress disorder (PTSD): A pilot study. Applied Psychophysiology Biofeedback, 36(1), 27–35. 10.1007/S10484-010-9141-Y/METRICS

Taulu, S., Simola, J., & Kajola, M. (2004). Clinical applications of the signal space separation method. International Congress Series, 1270(C), 32–37. 10.1016/J.ICS.2004.05.004

Treacher, A. H., Garg, P., Davenport, E., Godwin, R., Proskovec, A., Bezerra, L. G., Murugesan, G., Wagner, B., Whitlow, C. T., Stitzel, J. D., Maldjian, J. A., & Montillo, A. A. (2021). MEGnet: Automatic ICA-based artifact removal for MEG using spatiotemporal convolutional neural networks. NeuroImage, 241(June), 118402. 10.1016/j.neuroimage.2021.118402

Turcu, A. M., Ilie, A. C., Ştefăniu, R., Ţăranu, S. M., Sandu, I. A., Alexa-Stratulat, T., Pîslaru, A. I., & Alexa, I. D. (2023). The Impact of Heart Rate Variability Monitoring on Preventing Severe Cardiovascular Events. Diagnostics, 13(14). 10.3390/DIAGNOSTICS13142382

Usui, H., & Nishida, Y. (2017). The very low-frequency band of heart rate variability represents the slow recovery component after a mental stress task. PLoS ONE, 12(8). 10.1371/JOURNAL.PONE.0182611

van Driel, J., Olivers, C., & Fahrenfort, J. (2019). High-pass filtering artifacts in multivariate classification of neural time series data. BioRxiv, 530220. 10.1101/530220

Virtanen, P., Gommers, R., Oliphant, T. E., Haberland, M., Reddy, T., Cournapeau, D., Burovski, E., Peterson, P., Weckesser, W., Bright, J., van der Walt, S. J., Brett, M., Wilson, J., Millman, K. J., Mayorov, N., Nelson, A. R. J., Jones, E., Kern, R., Larson, E., … Vázquez-Baeza, Y. (2020). SciPy 1.0: fundamental algorithms for scientific computing in Python. Nature Methods, 17(3), 261–272. 10.1038/s41592-019-0686-2

Vrba, J., & Robinson, S. E. (1999). Functional neuroimaging by synthetic aperture magnetometry (SAM). Recent Advances in Biomagnetism, 2–5.

Vrba, J., & Robinson, S. E. (2001). Signal processing in magnetoencephalography. Methods, 25(2), 249–271. 10.1006/meth.2001.1238

Vrba, J., & Wilson, H. (2007). Comparison of external noise cancellation in MEG. International Congress Series, 1300, 603–606. 10.1016/J.ICS.2007.01.061

Wasimuddin, M., Elleithy, K., Abuzneid, A. S., Faezipour, M., & Abuzaghleh, O. (2020). Stages-based ECG signal analysis from traditional signal processing to machine learning approaches: A survey. IEEE Access, 8, 177782–177803. 10.1109/ACCESS.2020.3026968

Wolters, F. J., Segufa, R. A., Darweesh, S. K. L., Bos, D., Ikram, M. A., Sabayan, B., Hofman, A., & Sedaghat, S. (2018). Coronary heart disease, heart failure, and the risk of dementia: A systematic review and meta-analysis. In Alzheimer’s and Dementia (Vol. 14, Issue 11, pp. 1493–1504). 10.1016/j.jalz.2018.01.007

