## Supplemental File for "M-ECG: Extracting Heart Signals with a Novel Computational Analysis of Magnetoencephalography Data"

### Supplemental Figures

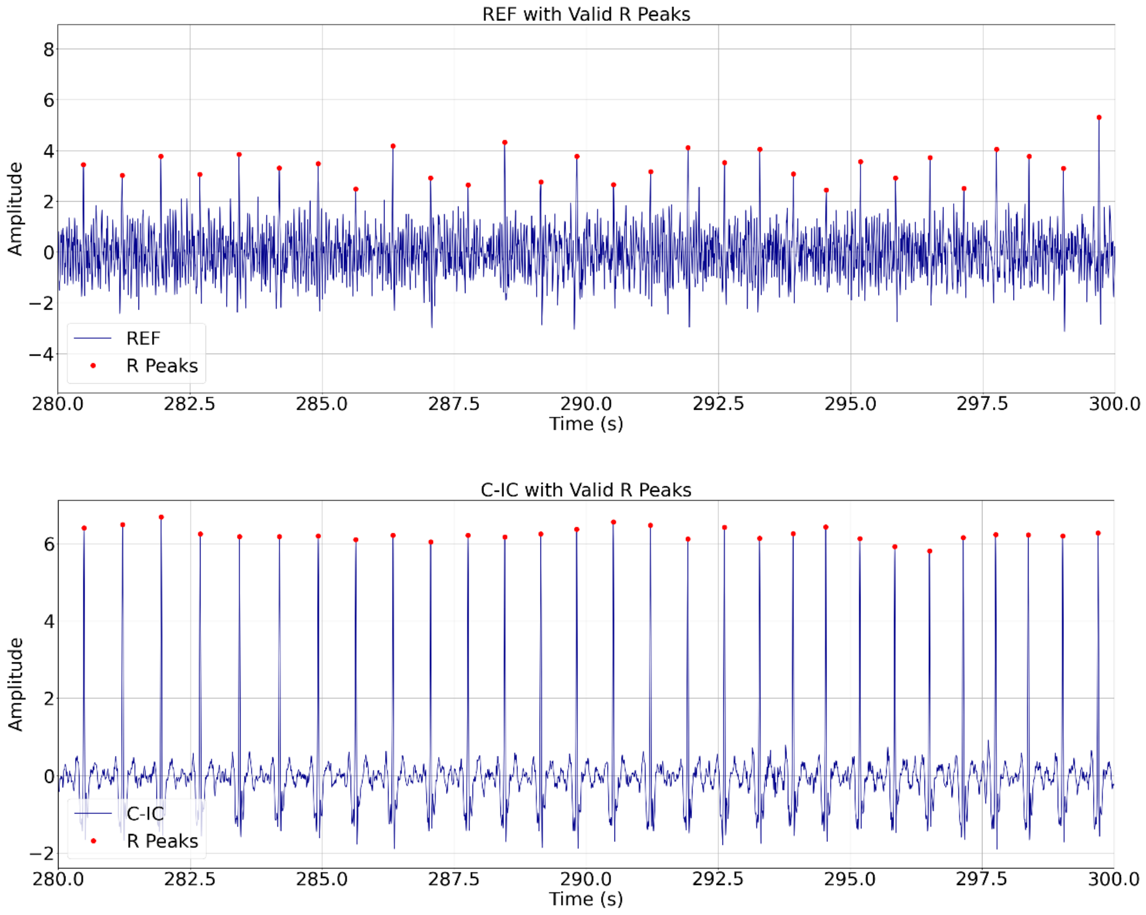


**Supplemental Fig. 1.** Automatically detected valid R peaks for REF and C-IC (280 – 300 s, participant 2).

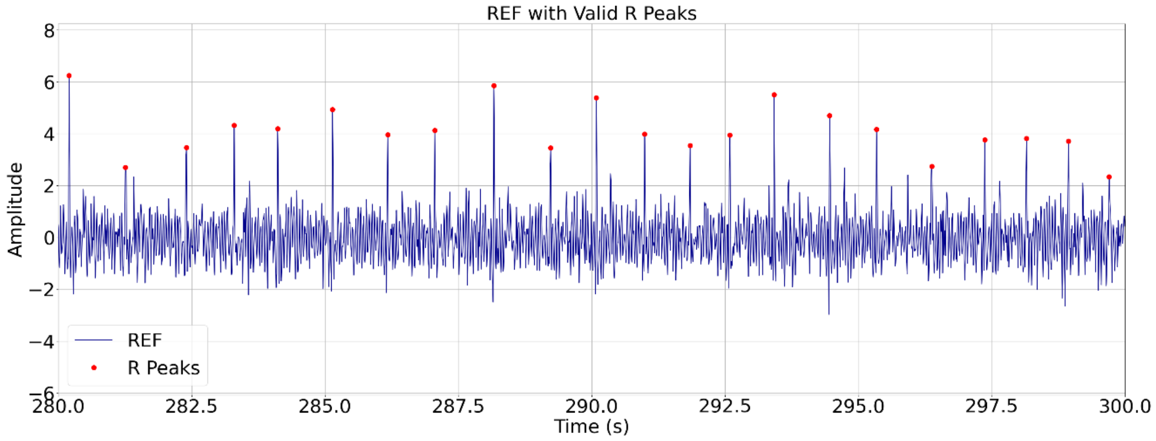

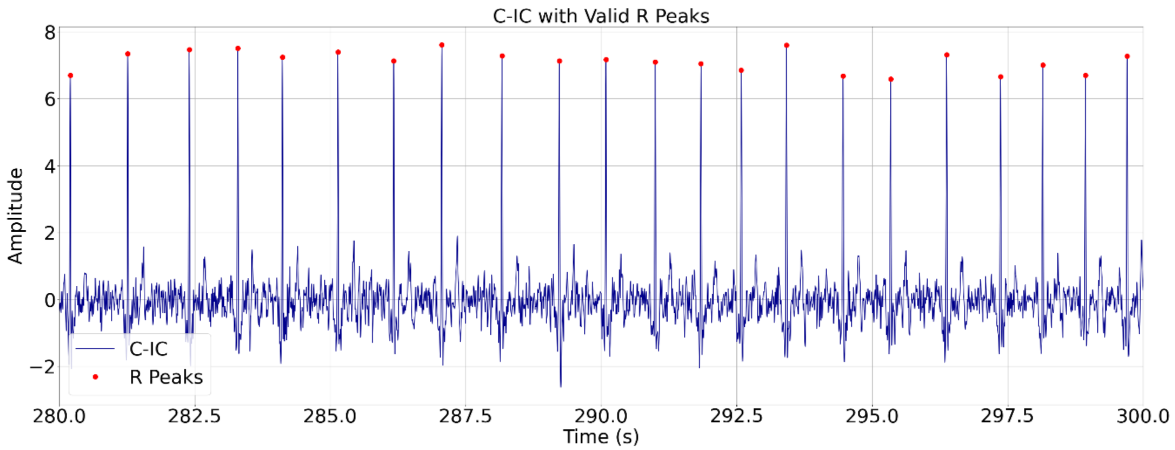


**Supplemental Fig. 2.** Automatically detected valid R peaks for REF and C-IC (280 – 300 s, participant 3).

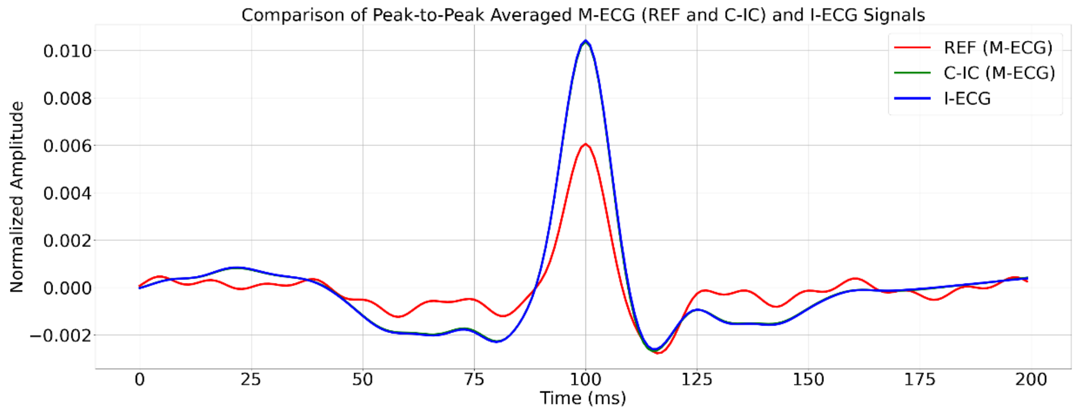


**Supplemental Fig. 3.** Comparison of averaged peak-to-peak REF, C-IC, and I-ECG signals with overlapping C-IC and I-ECG values (participant 2).

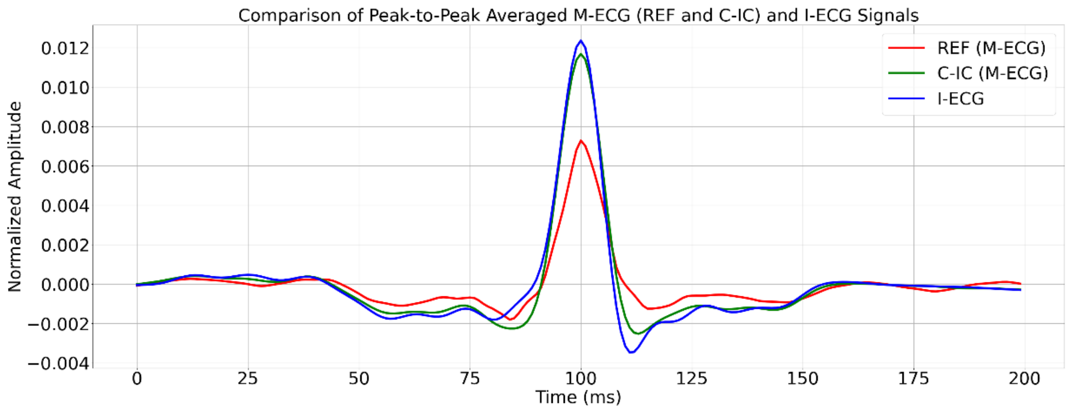


**Supplemental Fig. 4.** Comparison of averaged peak-to-peak REF, C-IC, and I-ECG signals (participant 3).

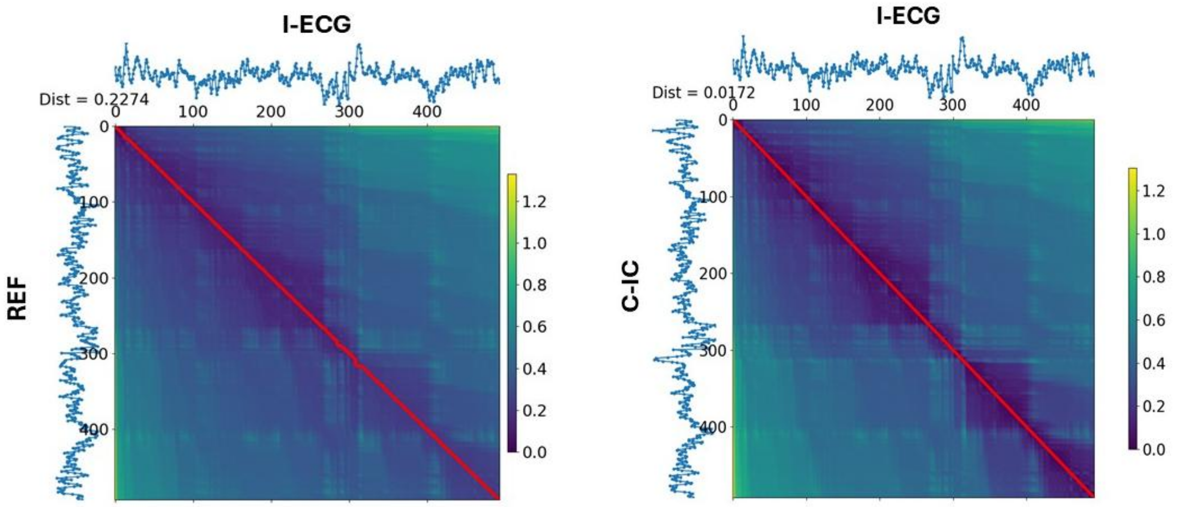


**Supplemental Fig. 5.** DTW of RR intervals (participant 2).

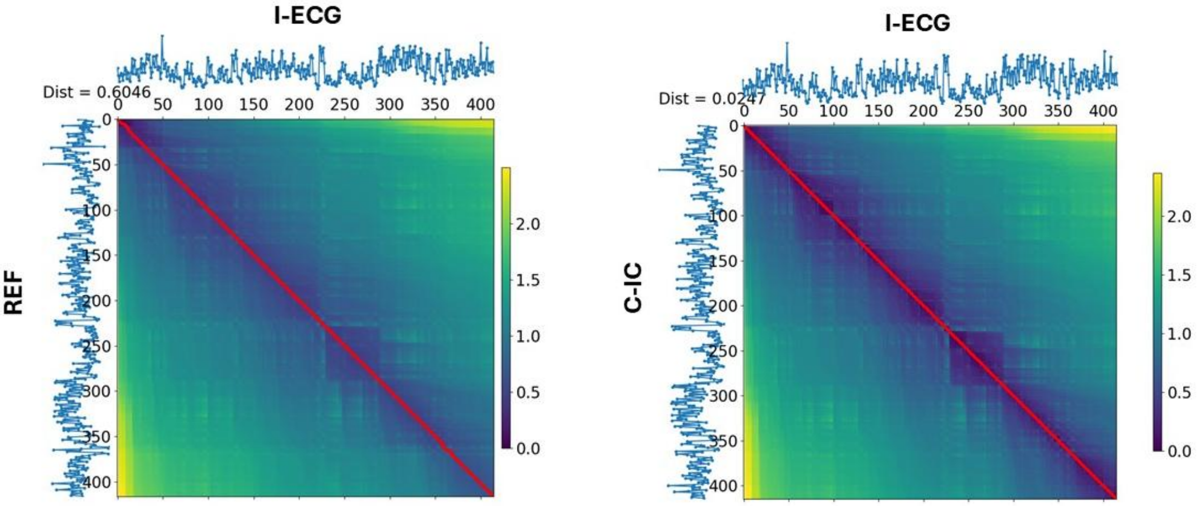


**Supplemental Fig. 6.** DTW of RR intervals (participant 3).

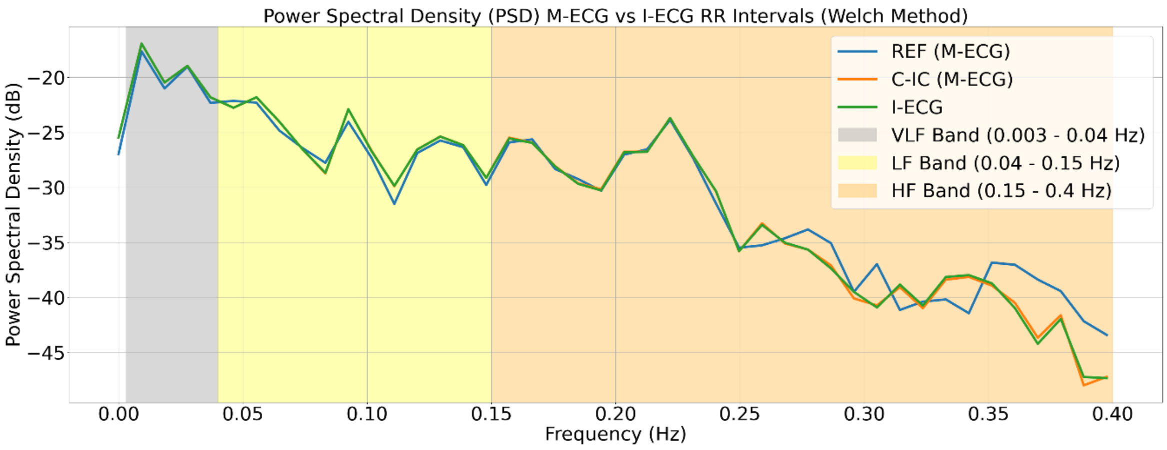


**Supplemental Fig. 7.** PSD between REF, C-IC, and I-ECG RR intervals (participant 2).

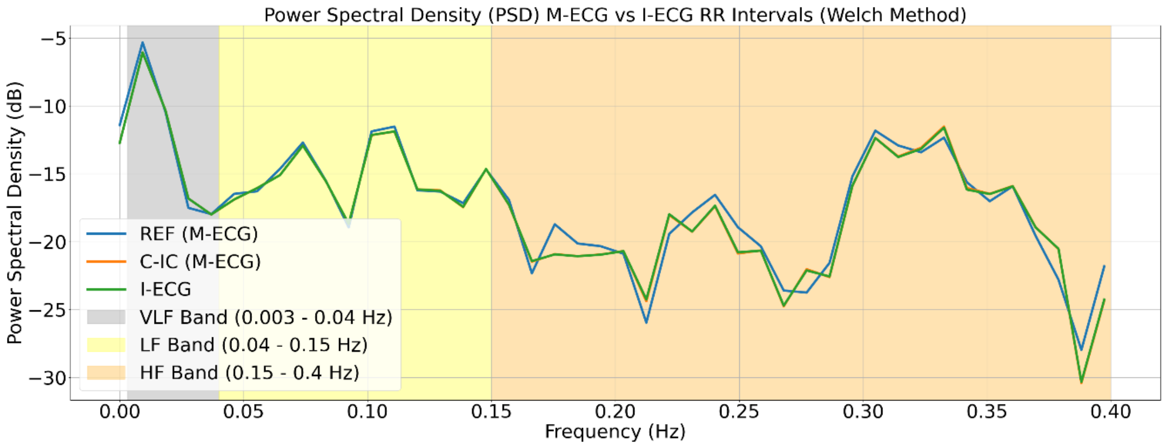


**Supplemental Fig. 8.** PSD between REF, C-IC, and I-ECG RR intervals (participant 3).

**Supplemental Table**

**Table 1**

Average processing time for M-ECG extraction for all 3 participants

| Participant | REF Processing Time (s) | C-IC Processing Time (s) |
|---|---|---|
| 1 | 2.79 | 11.56 |
| 2 | 25.25 | 36.07 |
| 3 | 33.24 | 36.6 |

Table 1 shows the average processing time for REF and C-IC extraction across three participants. C-IC processing times were consistently higher than REF, reflecting its increased computational load. The processing times were computed as the mean over ten execution runs for each participant and analysis type. The analysis was performed on a system equipped with an Intel Core i7-8700 processor (3.19 GHz), 32 GB of RAM, and a 64-bit Windows 11 operating system.
